# Left-handedness is associated with greater fighting success in humans

**DOI:** 10.1101/555912

**Authors:** Thomas Richardson, R. Tucker Gilman

**Affiliations:** School of Earth and Environmental Sciences, University of Manchester, United Kingdom

**Keywords:** Sexual-selection, Left-handedness, Fighting hypothesis, boxing, mixed martial arts

## Abstract

Left-handedness is a costly, sexually dimorphic trait found at low frequencies in all human populations. How the handedness polymorphism is maintained is unclear. The fighting hypothesis argues that left-handed men have a negative frequency-dependent advantage in violent intrasexual competition giving them a selective advantage. In support of this, many studies have found that left-handed men are overrepresented among modern professional fighters, but studies typically find no difference in fighting success between left and right-handed fighters. We studied over 13,800 professional boxers and mixed martial artists of varying abilities in three of the largest samples to test this hypothesis to date, finding robust evidence that left-handed fighters have greater fighting success. This held for both male and female fighters, and for both percentage of fights won and an objective measure of fighting ability. We replicated previous results showing that left-handed fighters are strongly overrepresented in professional combat sports, but left-handed fighters did not show greater variance in fighting ability, a hypothesis suggested in previous studies. Overall we find strong evidence consistent with the fighting hypothesis.

## Introduction

Left-handedness is a cross-culturally universal, heritable phenotype in humans [1] that is thought to be associated with fitness costs ([2,3], reviewed in [4], but see [5]). Typically around 11% of the population is left-handed [6] and though exact numbers vary with culture [7], left-handers are always a minority. Since left-handedness is under direct negative selection, its persistence in humans is an evolutionary puzzle.

One explanation for the persistence of left-handedness is the fighting hypothesis [8]. This argues that the polymorphism in human handedness is maintained due to a negative frequency-dependent advantage that left-handedness confers to males in combat (see [9] for theoretical support, and [10] for a review of empirical evidence as well as alternatives). According to this theory, right-handed males lack experience fighting rare left-handed males, while left-handed males accumulate plenty of experience fighting right-handed males, putting them at a selective advantage. Combined with the intrinsic fitness costs of left-handedness, this would explain the universal pattern of low but stable levels of left-handers in all studied populations. There is mounting evidence that intrasexual contest competition such as fighting has been a key component of sexual selection on human males ([11,12] reviewed in [13 and 14]). Modern males may possess adaptations to assist them in fighting and assessing opponents’ fighting ability [15]. Handedness could therefore be considered a sexually selected trait in males, and may be expressed in females a by-product [9].

Consistent with the fighting hypothesis, there is a wealth of evidence that left-handers are overrepresented in combat sports. Sports are particularly relevant systems for testing theories based on intrasexual competition, as they are thought to have evolved culturally as a display for males to advertise fighting and competitive ability [16]. Overrepresentation of left-handers has been seen in boxing [17–19], mixed martial arts or MMA [20–23], wrestling [24], Judo [25], and Karate and Taekwondo [26]. Left-handers are also overrepresented in many other sports, though crucially only sports requiring direct interaction with an opponent [27,28]. As they are rare, left-handers may gain an advantage because their actions are more difficult to predict [29–31], perhaps due to attentional biases towards the right hand of an opponent [32], which in combat sports is typically used for power strikes. If left-handed men are disproportionately successful in combat sports when they are rare, it is not unreasonable to assume they would also be successful in ancestral environments where physical violence and competition were likely much more common than today [14].

Studies of the fighting hypothesis in martial artists typically do not find that left-handed fighters are more likely to win fights (e.g., [20], but see [19]). However, previous studies have often used small sample sizes (e.g. [17]) or only assessed the very best members of a particular sport (e.g., [19, 24]). Any advantages are likely to be small as a large advantage would lead to an increase in the frequency of left-handed fighters until the advantage exactly offsets the costs of being left-handed, which may be small in populations with access to modern healthcare [2]. Thus, detecting the effect of left-handedness on fighting success may require very large sample sizes. Likewise, top fighters by definition have little variance in fight success, making detecting relationships in these datasets difficult. Top fighters may also have encountered enough left-handed opponents that any advantages due to unfamiliarity would be diminished. Evidence for whether left-handed fighters perform better than right-handed fighters is thus inconclusive. The present studies tested whether left-handed fighters are better than right-handed fighters in 3 large samples consisting of professional fighters at a variety of ability levels. In particular, one of our samples comprised the majority of boxers professionally active at the time of writing.

Previous studies also used win percentage records, number of wins, or ranking from a single tournament as proxies of fighting ability. These may fail to capture long term fighting performance, particularly for fighters with 0 losses, (which gives a win percentage of 1 regardless of the number of fights). These metrics also do not weight wins by quality of opponent, and fail to include how fighters beat their opponent. For example, winning a boxing match by having a better judges’ score after 10 rounds may indicate less physical dominance than a win by knockout in the first round. In our samples we excluded fighters who had few fights, and additionally compared left and right-handed boxers using their BoxRec score, a comprehensive measure of fighting ability that takes into account the type and swiftness of victory and opponent quality (see http://boxrec.com/media/index.php/BoxRec_Ratings_Description for a description of how a BoxRec score is calculated).

The fighting hypothesis for the evolution of left-handedness is based on male-male contest competition, but there is no reason to expect the frequency-dependent advantage of left-handedness in combat to be confined to males. However, there have been almost no of the success of left-handed female fighters. To remedy this, one of our samples consisted exclusively of female professional boxers and our sample of MMA fighters included women as well as men. Additionally, comparison of the left-hand advantage in male and female fighters allows us to investigate negative frequency-dependence. If there are fewer left-handed female fighters than male ones, the fighting hypothesis would predict left-handed female fighters would have a larger advantage.

Lastly, a previous study by Dochtermann et al. [22] demonstrated that left-handed MMA fighters show greater variance in probability of winning a fight than right-handed fighters. They argue that this is because the advantage left-handed fighters possess increases the probability that they will reach professional level compared to right-handers even if they are less skilled. We attempted to replicate this finding in our samples.

In summary, we investigated representation and fighting success of left-handers in 3 of the largest samples tested thus far, consisting of professional male and female boxers and MMA fighters of varying abilities. For boxers, we also tested the difference between left and right-handers in BoxRec scores, a holistic measure of fighting ability. Our study provides one of the most powerful tests of the fighting hypothesis attempted to date.

## Samples

Our first sample comprised every male professional boxer in the world listed as ‘active’ on www.boxrec.com at the time of writing (January 2019). BoxRec.com is a community-run boxing website that aims to document the careers of every professional boxer to have ever taken part in a recorded match. Boxers are listed as active if they have fought in an officially licensed bout in the past 12 months. Our second sample comprised all professional female boxers listed on www.boxrec.com for which stance data was available. For the female sample we included both active and retired boxers, as this ensured a large sample. Finally our third sample comprised all the MMA fighters listed on ufcstats.com at the time of writing. ufcstats.com is a comprehensive, respected MMA database that is the official statistics provider to the Ultimate Fighting Championship (UFC).

We excluded fighters with fewer than 5 fights, as their fight record is too preliminary to accurately reflect their fighting ability. We additionally excluded fighters with a win percentage of 20% of less. Many of these fighters are likely what are referred to in boxing slang as “tomato cans”: uncompetitive fighters who take matches with opponents they have little chance of beating simply to earn money. They are often matched against young up-and-coming fighters in order to gain the fighter more wins on their record. For these reasons their win percentage and Boxrec score may not reflect their fighting ability, and as such they were excluded.

The final samples consisted of 10445 male boxers, (8666 right-handed and 1779 left-handed), 1314 female boxers, (1150 right-handed and 164 left-handed fighters) and 2100 MMA fighters (1707 right-handed and 393 left-handed fighters).

## Results

All statistics were run in R [34], and all data and analysis code is available on the open science foundation (https://osf.io/x3unr/). For all samples, the number of fights left- and right-handed fighters had participated in, fighter ages, win percentages and BoxRec scores were all non-normally distributed, so nonparametric statistics were used throughout.

A Mann-Whitney U test showed that left-handed male boxers did not differ in age (*p* = 0.88) from right-handed boxers. For female fighters, age was not analysed as some boxers were retired, deceased or not currently active. Age was not available for the MMA fighters. Mann-Whitney U tests found no significant differences in number of fights between left- and right-handed fighters among male boxers (*p* = 0.40) and female boxers (*p* = 0.69) though left-handed MMA fighters did have marginally more fights than right-handed fighters (*p* = 0.047). Additionally, t-tests showed that left- and right-handed MMA fighters did not differ in overall weight, height or arm length (also known as “reach”) (all *p* > 0.19). This data was not available for either sample of boxers.

### Are left-handers overrepresented among professional fighters?

To test whether left-handed fighters were overrepresented in our samples we ran three separate, one-tailed binomial tests against percentages of left-handers found in a large representative, western population [6]. We tested the percentage of left-handed male boxers against the percentage of left-handed men (12.6%) and female boxers against the percentage of left-handed women (9.9%) in the general population. The MMA sample included both male and female fighters, so was tested against the percentage of left-handed men, as this was the most conservative test of our hypothesis. Table 1 shows that left-handed fighters were significantly overrepresented in all three samples (all *p* ≤ 0.001).

**Table 1.** results of Binomial tests of % of left-handed fighters against % of left-handed people in the general population

| the general population |  |  |  |
| --- | --- | --- | --- |
| Sample | % left-handed fighters in sample | % left-handers in general population | p-value |
| Male boxers | 17.0 | 12.6 | < 0.0001 |
| Female boxers | 12.5 | 9.9 | = 0.001 |
| MMA fighters | 18.7 | 12.6 | < 0.0001 |

### Do left-handed fighters possess greater fighting ability than right-handed fighters?

We compared the fighting success of left- and right-handed fighters with one-tailed Mann-Whitney U tests. Each of the 3 samples was compared separately by win percentages, and the samples of male and female boxers were also compared by BoxRec scores. We calculated the measure of stochastic superiority [35,36] as an effect size for each comparison. The measure of stochastic superiority is the probability that a randomly selected left-handed fighter would have a higher win percentage/BoxRec score than a randomly selected right-handed fighter.

Among male boxers, the probability that a randomly selected left-handed fighter would have a higher BoxRec score than a randomly selected right-handed fighter was 52.4%, which a Mann-Whitney test showed was significant (*p*=0.00069). The measure of stochastic superiority for win percentage was also 52.4%, which was also significant (*p*=0.0007). Thus left-handed male boxers have significantly higher BoxRec scores and win percentages than right-handed male boxers.

Among female boxers, the probability that randomly selected left-hander showed a higher BoxRec score was 53.9%, which a Mann-Whitney test showed was marginally significant (*p*=0.053). The measure of stochastic superiority for win percentage was 54.5%, which was significant (*p*=0.031). Thus left-handed female boxers showed significantly higher BoxRec scores but not win percentages.

Among MMA fighters, the probability that a randomly sampled left-handed fighter showed a higher win percentage than a randomly selected right-handed fighter was 53.5%, which was significant (*p*=0.016). A Spearman’s correlation between total number of fights and win percentage revealed a negative correlation (*r* = −0.14, p<0.0001), indicating that this effect is not due to left-handed fighters having more fights. Thus left-handed MMA fighters showed significantly higher win percentages than right-handed MMA fighters.

### Do left-handed fighters show greater variance than right-handed fighters?

We compared the variance in BoxRec scores and win percentages among left- and right-handers by bootstrapping differences in variance (10,000 samples), with bias correction and acceleration following [37] to obtain robust p-values. All p-values are one-tailed. Left-handed male boxers showed higher variance in BoxRec scores (*p* = 0.002) but not in win percentages (*p* = 0.74). Left-handed female fighters did not differ from right-handed female fighters in the variance of their BoxRec scores (*p* = 0.80) or win percentages (*p* = 0.77). Likewise left-handed MMA fighters did not differ from right-handed MMA fighters in the variance of their win percentages (*p* = 0.54).

### Does the left-hand advantage show negative frequency-dependence?

The prevalence of left-handedness in female boxers was much lower than in male boxers (17% vs 12.5%), while the magnitude of left-hand advantage in the BoxRec scores of female fighters was higher (54.5% vs 52.5%). If the advantage left-handed fighters have is negative frequency-dependent, then we might expect left-handed female boxers to have a relatively larger advantage than left-handed male boxers. To investigate this, we compared the measures of stochastic superiority in the BoxRec scores of male and female boxers, and we bootstrapped a confidence interval around the difference (10,000 samples). The difference in the advantage of left-handed female and male boxers was not significantly different from 0 (bias corrected, accelerated p-value = 0.29).Thus, we have no evidence that female boxers experience a greater left-hand advantage than male boxers.

## Discussion

Across three samples, we found that left-handed boxers and MMA fighters are both overrepresented in their respective sports and are more successful fighters. In male boxers, these effects held for both win percentages and BoxRec scores, where BoxRec scores are a more comprehensive measure of boxing ability. In female boxers we found that left-handed fighters showed higher BoxRec scores but not higher win percentages. Our results are consistent with the fighting hypothesis that left-handedness is maintained in populations because it provides a advantage in contest competition.

Our finding that left-handed fighters have better records than right-handed fighters in both male boxers and MMA fighters contrasts to most previous studies (e.g. [18,20,21], but see [19]). Two factors may have played a role. Firstly, the effect is small and may only be detectable in large samples such as ours. Second, it may not be detectable in datasets with low variance in fighting ability, such as when studies use samples of only elite fighters (e.g., [18]). The fact that we find similar results in both win percentages and BoxRec scores, which are a more complete measure of boxing ability, lead us to believe our results are robust.

Our positive finding for MMA fighters may be surprising, as a similar study [21] did not find a significant advantage of left-handedness in a sample approximately 70% of the size of ours. The study collected data from the same website we did ~6 years earlier, so its data set likely overlaps with ours. The different results may be due to the choice of analyses, or to the fact that the study did not exclude fighters with few fights as we did. It is noteworthy that in [21], left-handed fighters had a non-significantly higher win percentage, so the trend reported is consistent with our results.

We found that left-handed female boxers showed better BoxRec scores than right-handed female boxers. As there were fewer left-handed fighters in the female sample than the male sample (12.6% to male’s 17.3%), we tested whether the left-hand advantage seen in female fighters was higher than that of male fighters. Left-handed female fighters being less numerous and having greater success than their male counterparts would be consistent with the fighting hypothesis, in that it suggests a negative frequency-dependent advantage. However we did not find this. That the left-handed advantage in combat is negative frequency-dependent remains to be convincingly demonstrated, and is a crucial topic of future research. This might be investigated by comparing fighting leagues with varying levels of left-handers, or by testing whether increased contact with left-handed opponents over a fighter’s career increases his/her probability of winning.

Unlike Dochtermann et al. [22], overall we found little evidence that left-handed fighters showed higher variance in fighting ability. Across all samples, only male left-handed boxers showed significantly higher variance, and then only in BoxRec scores. The difference in results could be attributed to the fact that Dochtermann et al. tested variance in the probability of a fighter to win a single given fight, whereas we examined variance in fighting success as measured by a fighter’s record over their career thus far. It is possible that coaches (many of whom may suspect the existence of a left-handed advantage) or the left-handed fighters themselves adapt their training to compensate for their fighter’s lower skills. However we warn that cross sectional data, such as ours and that of Dochtermann et al., are limited in their ability to answer this question. Longitudinal work that tracks whether left-handed amateurs are more likely to reach professional level regardless of initial skill would be valuable, and shed more light on this interesting hypothesis.

## Conclusion

In conclusion, we present strong evidence that left-handed fighters show greater fighting success, consistent with the fighting hypothesis. Our study also provides further evidence that left-handed fighters are overrepresented in combat sports. We demonstrate these effects in 3 of the largest samples to test the hypothesis to date, using both male and female fighters, and using multiple measures of fighting competence. Future research linking fighting stance to fitness costs associated with handedness, as well as more direct work investigating the negative frequency-dependent nature of the left-hand advantage, is required.

**Figure 1:**
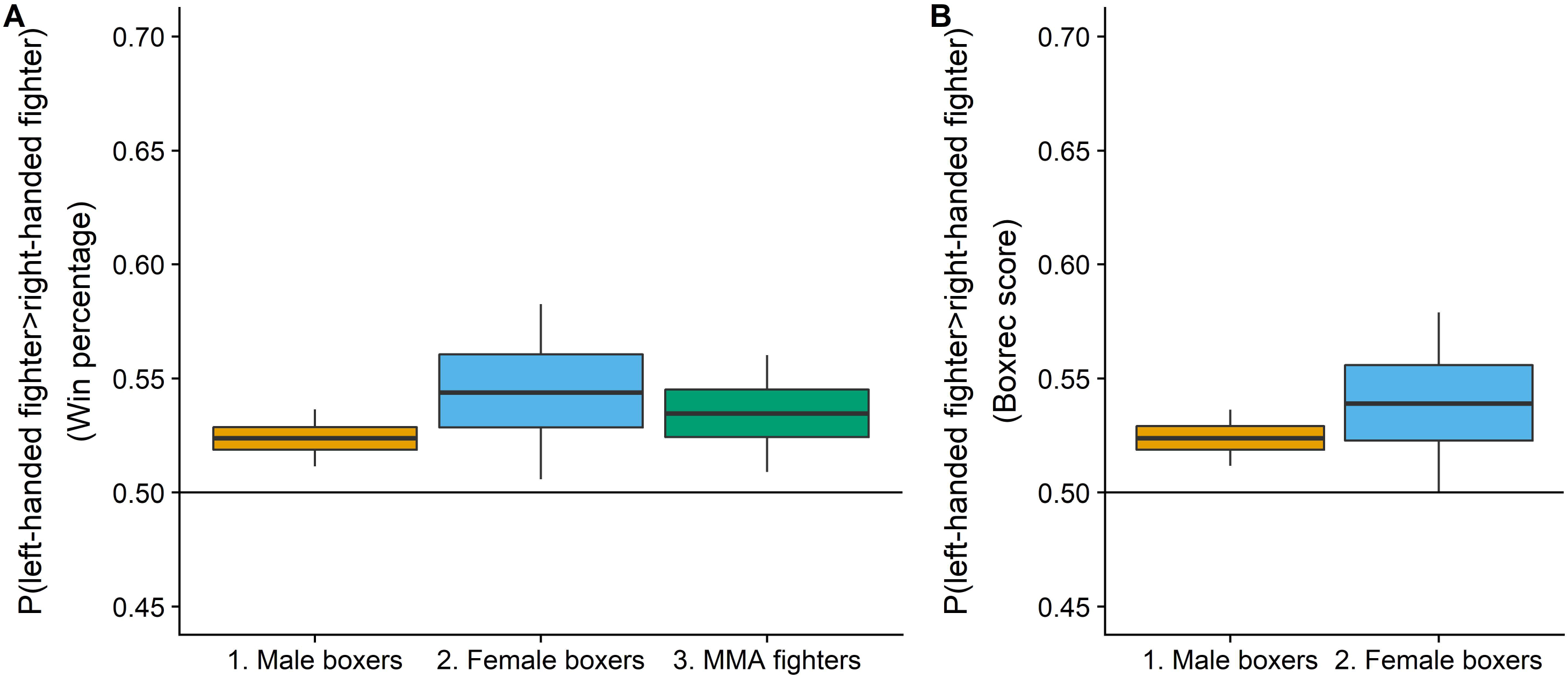
The probability that a randomly selected left-hander showed a higher (A) win percentage and (B) BoxRec score than a randomly selected right-hander. Boxes indicate 50% and whiskers indicate 90% bootstrapped confidence intervals (5000 samples).

## Acknowledgements

We thank Rebecca Lewis and the other members of the Gilman lab for critical comments on the draft of this paper as well as the community of BoxRec.com and James Carr for insight into the world of boxing.

## Data, code and materials

All data associated with this manuscript, as well as R code to conduct statistical analyses and create the graphs are uploaded as part of the supplementary material and can be additionally found at https://osf.io/x3unr/.

## Author’s contributions

TR conceived of and designed the study, obtained the data and wrote the manuscript. Both TR and RTG carried out statistical analysis. RTG critically revised the manuscript.

## References

1. Medland SE, Duffy DL, Wright MJ, Geffen GM, Hay DA, Levy F, Van-Beijsterveldt CE, Willemsen G, Townsend GC, White V, Hewitt AW. Genetic influences on handedness: data from 25,732 Australian and Dutch twin families. Neuropsychologia. 2009 Jan 1;47(2):330–7

2. Coren S, Halpern DF. Left-handedness: a marker for decreased survival fitness. Psychological bulletin. 1991 Jan;109(1):90

3. Dragovic M, Hammond G. Handedness in schizophrenia: a quantitative review of evidence. Acta Psychiatrica Scandinavica. 2005 Jun;111(6):410–9

4. Llaurens V, Raymond M, Faurie C. Why are some people left-handed? An evolutionary perspective. Philosophical Transactions of the Royal Society B: Biological Sciences. 2008 Dec 5;364(1519):881–94

5. Zickert N, Geuze RH, van der Feen FE, Groothuis TG. Fitness costs and benefits associated with hand preference in humans: A large internet study in a Dutch sample. Evolution and Human Behavior. 2018 Mar 1;39(2):235–48

6. Gilbert AN, Wysocki CJ. Hand preference and age in the United States. Neuropsychologia. 1992 Jul 1;30(7):601–8

7. Raymond M, Pontier D. Is there geographical variation in human handedness?. Laterality: Asymmetries of Body, Brain and Cognition. 2004 Jan 1;9(1):35–51

8. Raymond M, Pontier D, Dufour AB, Møller AP. Frequency-dependent maintenance of left handedness in humans. Proc. R. Soc. Lond. B. 1996 Dec 22;263(1377):1627–33

9. Billiard S, Faurie C, Raymond M. Maintenance of handedness polymorphism in humans: a frequency-dependent selection model. Journal of Theoretical Biology. 2005 Jul 7;235(1):85–93

10. Groothuis TG, McManus IC, Schaafsma SM, Geuze RH. The fighting hypothesis in combat: how well does the fighting hypothesis explain human left-handed minorities?. Annals of the New York Academy of Sciences. 2013 Jun 1;1288(1):100–9

11. Hill AK, Hunt J, Welling LL, Cárdenas RA, Rotella MA, Wheatley JR, Dawood K, Shriver MD, Puts DA. Quantifying the strength and form of sexual selection on men’s traits. Evolution and Human Behavior. 2013 Sep 1;34(5):334–41

12. Kordsmeyer TL, Hunt J, Puts DA, Ostner J, Penke L. The relative importance of intra-and intersexual selection on human male sexually dimorphic traits. Evolution and Human Behavior. 2018 Jul 1;39(4):424–36

13. Puts DA, Jones BC, DeBruine LM. Sexual selection on human faces and voices. Journal of sex research. 2012 Mar 1;49(2–3):227–43

14. Puts DA. Beauty and the beast: Mechanisms of sexual selection in humans. Evolution and Human Behavior. 2010 May 1;31(3):157–75

15. Třebický V, Stirrat M, Havlíček J. Fighting Assessment. In: T. K. Shackelford, V. A. Weekes-Shackelford (eds.), Encyclopedia of Evolutionary Psychological Science Switzerland: Springer Nature p. 1–11

16. Deaner RO, Balish SM, Lombardo MP. Sex differences in sports interest and motivation: An evolutionary perspective. Evolutionary Behavioral Sciences. 2016 Apr;10(2):73

17. Gursoy R. Effects of left-or right-hand preference on the success of boxers in Turkey. British Journal of Sports Medicine. 2009 Feb 1;43(2): 142–4

18. Sorokowski P, Sabiniewicz A, Wacewicz S. The influence of the boxing stance on performance in professional boxers. AnthropologicAl review. 2014 Dec 1;77(3):347–53

19. Loffing F, Hagemann N. Pushing through evolution? Incidence and fight records of left-oriented fighters in professional boxing history. Laterality: Asymmetries of Body, Brain and []Cognition. 2015 May 4;20(3):270–86

20. Pollet TV, Stulp G, Groothuis TG. Born to win? Testing the fighting hypothesis in realistic fights: left-handedness in the Ultimate Fighting Championship. Animal behaviour. 2013 Oct 1;86(4):839–43

21. Baker J, Schorer J. The Southpaw Advantage?-Lateral Preference in Mixed Martial Arts. Plos One. 2013 Nov 19;8(11):e79793

22. Dochtermann NA, Gienger CM, Zappettini S. Born to win? Maybe, but perhaps only against inferior competition. Animal Behaviour. 2014(96):e1–3

23. Pollet TV, Riegman BR. Opponent left-handedness does not affect fight outcomes for Ultimate Fighting Championship hall of famers. Frontiers in psychology. 2014 Apr 30;5:375

24. Ziyagil MA, Gursoy R, Dane Ş, Yuksel R. Left-handed wrestlers are more successful. Perceptual and motor skills. 2010 Aug;111(1):65–70

25. Tirp J, Baker J, Weigelt M, Schorer J. Combat stance in judo–Laterality differences between and within competition levels. International Journal of Performance Analysis in Sport. 2014 Apr 1;14(1):217–24

26. Cingoz YE, Gursoy R, Ozan M, Hazar K, Dalli M. Research on the Relation between Hand Preference and Success in Karate and Taekwondo Sports with Regards to Gender. Advances in Physical Education. 2018 Aug 1;8(03):308

27. Aggleton JP, Wood CJ. Is there a left-handed advantage in “ballistic” sports?. International Journal of Sport Psychology. 1990 Jan

28. Grouios G, Tsorbatzoudis H, Alexandris K, Barkoukis V. Do left-handed competitors have an innate superiority in sports?. Perceptual and motor skills. 2000 Jun;90(3_suppl):1273–82

29. Hagemann N. The advantage of being left-handed in interactive sports. Attention, Perception, & Psychophysics. 2009 Oct 1;71(7):1641–8

30. Loffing F, Schorer J, Hagemann N, Baker J. On the advantage of being left-handed in volleyball: further evidence of the specificity of skilled visual perception. Attention, Perception, & Psychophysics. 2012 Feb 1;74(2):446–53

31. Loffing F, Hagemann N, Schorer J, Baker J. Skilled players’ and novices’ difficulty anticipating left-vs. right-handed opponents’ action intentions varies across different points in time. Human movement science. 2015 Apr 1;40:410–21

32. Marzoli D, Lucafò C, Pagliara A, Cappuccio R, Brancucci A, Tommasi L. Both right-and left-handers show a bias to attend others’ right arm. Experimental brain research. 2015 Feb 1;233(2):415–24

33. Loffing F, Hagemann N. Pushing through evolution? Incidence and fight records of left-oriented fighters in professional boxing history. Laterality: Asymmetries of Body, Brain and Cognition. 2015 May 4;20(3):270–86

34. Team RC. R: A language and environment for statistical computing

35. McGraw KO, Wong SP. A common language effect size statistic. Psychological bulletin. 1992 Mar;111(2):361

36. Vargha A, Delaney HD. A critique and improvement of the CL common language effect size statistics of McGraw and Wong. Journal of Educational and Behavioral Statistics. 2000 Jun;25(2):101–32

37. DiCiccio TJ, Efron B. Bootstrap confidence intervals. Statistical science. 1996 Aug 1:189–212

